# Balance of AKT isoforms in intestinal barrier function. Implications for inflammatory bowel disease therapy

**DOI:** 10.1101/2023.07.04.547629

**Authors:** Teresa García-Prieto, Antonio Barbachano, Natalia Cuesta, María Dolores Sánchez, Alicia Arranz, Manuel Fresno

**Author notes:** In memoriam (Alicia Arranz).

## Abstract

WORKING MODEL

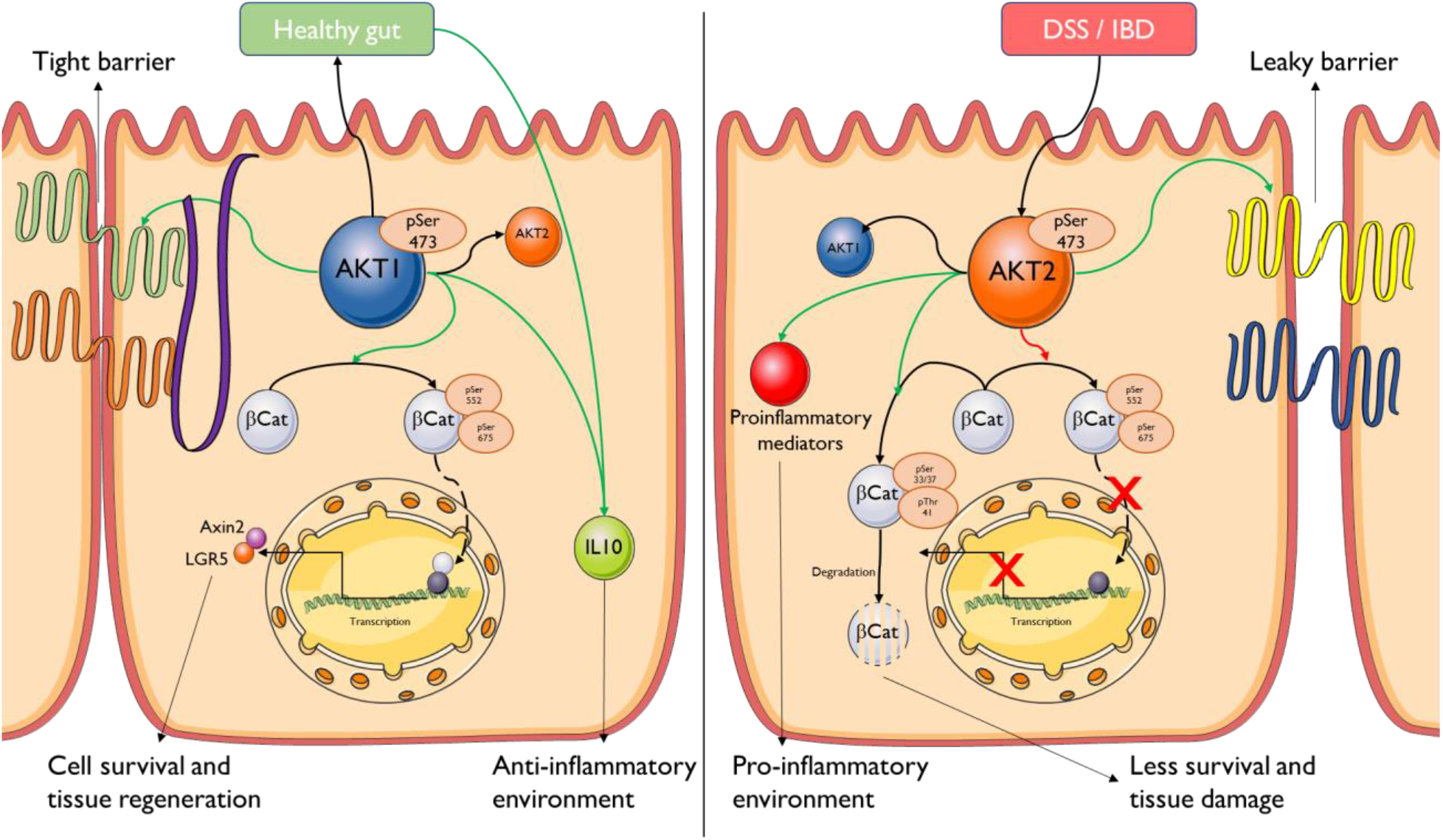

The leakiness of the intestinal epithelial barrier plays a major role in the development of inflammatory bowel disease (IBD). We show that either Akt1 overexpression or Akt2 chemical inhibition have similar effects on intestinal epithelial cells, enhancing the permeability of the epithelial barrier and affecting the expression and/or intracellular localization of the tight junctions’ proteins (TJP) whereas Akt1 inhibition or Akt2 overexpression show the opposite results. Akt1 overexpression or Akt2 inhibition also promotes activation of β-Catenin and Lgr5. Moreover, mouse intestinal organoids treated ex vivo with Akt1 inhibitors present dissembled and disorganized structures that mimic IBD histological features. Importantly, in DSS-induced IBD in mice, Akt2 inhibition strongly ameliorates disease with animals presenting healthy-like crypts, having tightened colon epithelial cells expressing TJP, and presenting an anti-inflammatory M2 macrophage phenotype. In contrast, treatment with Akt1 inhibitors enhances histological damage, TJP disorganization reduces Lgr5+ expression and promotes a more proinflammatory milieu further favoring the development of the disease. Those results suggest that a balance between Akt1 and Akt2 regulates the functionality of the intestinal epithelial barrier, cell renewal and inflammation and that Akt2 inhibition can be considered as a new therapy for IBD.

**THE PAPER EXPLAINED:** Inflammatory bowel disease (IBD), that includes Crohńs disease and ulcerative colitis, needs better therapeutic options. Importantly, IBD is a risk factor for the development of colorectal cancer. IBD is associated with epithelial barrier dysfunction, decreasing its permeability, and finally leading to chronic inflammation and disease. We show that the Akt1 and Akt2 protein kinase isoforms play different and opposite roles in the integrity of intestinal epithelial barrier by affecting the expression and/or the intracellular localization of tight junction proteins (TJP) responsible for barrier formation and reducing anti-barrier ones when Akt1 is overexpressed or Akt2 is inhibited. Those results were confirmed in ex vivo organoids, a system that mimics intestinal tissue organization, where Akt1 inhibition resulted in dissembled and disorganized structures that resembled those found in IBD patients. There is a balance between Akt1 and Akt2 activation on the epithelial cells, and when Akt1 is less abundant than Akt2, the barrier is disrupted by the dysregulation of the tightness of the barrier by the TJPs, leading also to an acute phase of inflammation. Epithelial cells are also incapable of renewing themselves by the absence of active Akt1, making the damage more severe, creating a pro-inflammatory niche that derives on chronic inflammation and severe IBD. Interestingly, those in vitro and ex vivo results can be translated to the whole animals. IBD disease and colon inflammation and histological alterations induced in mice by an irritant DSS could be either worsened with an Akt1 inhibitor or much improved with treatment with an Akt2 inhibitor, respectively. The inhibition of Akt2 may be considered as a treatment or co-treatment of IBD by regulating this balance to an Akt1 predominance environment. Thus, the use of Akt2 inhibitors, already developed, could be an alternative way to treat this disease.

## INTRODUCTION

Inflammatory bowel disease (IBD) comprises a group of diseases, including Crohńs disease (CD) and Ulcerative Colitis (UC), characterized by an hyperactivated and uncontrolled state of inflammation in the intestinal tract, involving intestinal epithelium damage and leading to epithelial barrier leakiness (Schulzke et al. 2009; Krug y Fromm 2020). There are multiple factors, not mutually exclusive, underlying to the development of IBD. Those include genetics factors (Gazouli et al. 2019), as it is already known that mutations on some innate immune receptors, such as TLR2, TLR4 and NOD2 may predispose to disease (Cheng et al. 2015; Hugot et al. 2001; Ogura et al. 2001). Composition of intestinal microbiota (Matsuoka y Kanai 2015) and external factors such as drugs, smoke or the diet may also affect (de Souza y Fiocchi 2016; Wallace 2014; Y.-Z. Zhang 2014). Moreover, IBD is a risk factor in the development of colorectal cancer, which is one of the most worrying sequelae of the disease. Therapies for IBD include blocking antibodies to inflammatory cytokines like TNF and treatments with immunomodulators in order to control the inflammation, but they may have secondary effects as a consequence of the depression of the immune system. A better understanding of the signaling leading to inflammation and their control represents a promising approach for therapy (Geremia et al. 2014).

The serine-threonine protein kinase Akt, also known as protein kinase B (PKB), is a master protein kinase in mammals, which has a role in important pathways such as glucose metabolism, cell proliferation and apoptosis regulation, transcription, and cell migration (Sheng et al. 2003; G. Lee et al. 2010; Whiteman, Cho, y Birnbaum 2002; Shiojima y Walsh 2002). Akt has three isoforms which are highly homologous. Although those 3 isoforms may play some redundant roles, it has been described that they present different functions too. Thus, Akt3 (PKBγ) has been implicated in neuronal development (T. Zhang et al. 2021; DuBois et al. 2019), Akt2 (PKBβ) in the glucose homeostasis and metabolism (Hay 2011; M. Chen et al. 2020; Sakamoto et al. 2006) and Akt1 (PKBα) in cell growth and angiogenesis (Somanath et al. 2006; Dummler et al. 2006; M. Y. Lee et al. 2014; Duggal et al. 2018; Héron-Milhavet et al. 2006).

In recent years, there are some evidences linking the development of IBD to the expression of Akt isoforms (T. Chen et al. 2017; Gazouli et al. 2019). More interestingly, mice lacking *Akt2* presented higher resistance to the development of the disease in response to DSS treatment, the chemical model of Ulcerative Colitis (Arranz et al. 2012). This effect was ascribed to polarization of macrophages to M2 in absence of *Akt2*, since those Akt2 KO mice presented lower levels of proinflammatory cytokines (TNFα, IL6, NO) and an increase in Arginase 1, a well-known marker of M2 macrophages. In contrast Akt1 KO mice were more susceptible and presented the opposite characteristics.

Intestinal epithelial cells (IECs) are responsible for several functions on the gut: nutrients absorbance, epithelial barrier maintenance, microbiota tolerance and pathogens response, innate immune system development and immune signaling. IECs express Akt1 and Akt2 in high levels but not Akt3 (Dufour et al. 2004). Akt1 promotes the survival of the IECs (H. M. Zhang et al. 2004) and Akt2 is implicated in the absorption and regulates NHE3, a Na+/H+ channel (Shiue et al. 2005). IECs are responsible for the maintenance of the intestinal barrier and have an important role in the development of innate immune response and intestinal inflammatory pathologies (J. Lee et al. 2018; Chelakkot, Ghim, and Ryu 2018; Schulzke et al. 2009). The integrity of this barrier is relevant for the intestinal function, mainly due to the expression and function of TJP (S. H. Lee 2015) present the intestinal epithelial cells, which modulate the permeability of the intestinal barrier in response to both physiological and pathological stimuli which modulate the permeability of the intestinal barrier in response to both physiological and pathological stimuli (Peterson and Artis 2014). Depending on the TJP, the epithelium would be more or less permeable to all kind of particles from viruses and bacteria to different molecules that can signal to both pro and anti-inflammatory state. Those proteins are key in the maintenance of the intestinal barrier (Weber et al. 2008) and there is an increasing interest on their potential role as biomarkers due to differences observed on CD and UC patients (Das et al. 2012) and on signal transduction (Takano et al. 2014; González-Mariscal, Tapia, and Chamorro 2008). Thus, restoring the epithelial barrier function may represent a promising therapeutic approach for IBD, but no therapies are currently available (Odenwald and Turner 2017).

Another important molecule implicated in the junctions between the intestinal cells is β-Catenin, that together with E-Cadherin, is one of the main actors in the adherents junction, and defects on this complex are implicated on IBD development (Mehta S et al. 2015). When β-Catenin is not complexed with E-Cadherin at the cell surface, it may signal through the WNT/β-Catenin complex entering into the nucleus where it can regulate the expression of a large variety genes of including other transcription factors implicated in cell maintenance and survival, such as Axin-2 or c-Myc, and other relevant proteins as LGR5, implicated in cell maintenance and survival which is also the main marker of intestinal stem cells (ISCs). (Chopra et al. 2010; Nusse y Clevers 2017; González-Sancho, Larriba, y Muñoz 2020). ISCs in healthy conditions, remain at the edge of the intestinal crypt, and can give rise to each one of the cell types that can be found on the crypt: epithelial, globet, Paneth, tuft cells, etc., but they may also differ in response to damage. Dysregulation of this pathway is also very important in the maintenance of the self-renewal of intestinal epithelium being one of the main drivers of colon carcinogenesis.

IECs can produce and respond to both pro and anti-inflammatory signals. The switch between these two stages is important in the development of IBD. Disruption of the intestinal epithelial barrier facilitates the entry of microbes into the intestinal wall and the subsequent activation of inflammation (Neurath 2014): Production of pro-inflammatory mediators and activation of M1 macrophages, followed by a failure on the regulation system by Treg lymphocytes and production of IL10 to finish the immune response, which leads to the uncontrolled and hyperactivated inflammation. In this regard, IL10 KO mice develop spontaneous colitis, (Shouval et al. 2014) pointing to the relevant role of the cytokine milieu.

Here we have addressed the role of Akt isoforms on IBD, both in vitro and in vivo. We found that Akt1 and Akt2 play an opposite role in IBD by differentially controlling TJP and IEC permeability as well as inflammation both in vitro, ex vivo and in vivo in animal models of IBD. We found that the use of an Akt2 inhibitor strongly improved DSS-induced IBD in mice and may represent a promising therapy.

## MATERIALS AND METHODS

### AKT STABLE OVEREXPRESSING CELLS

Four IECs were used; Caco2 (ATCC® HTB37™) and HT29 (ATCC® HTB38™) that are epithelial cells derived from human colorectal adenocarcinoma, SW480 (ATCC® CCL228™) from Dukes’ type B, colorectal adenocarcinoma and HCT116 (ATCC® CCL247™) derived from human colorectal carcinoma. IECs were cultured in MEM (Minimum Essential Medium Eagle) with 10% FBS (Fetal Bovine Serum) for Caco2 and 5% FBS for the rest of the cells and complemented with Streptomycin at 100 µg/mL and Penicillin 100 U/mL, Glutamine at 2 mM, NEAA and Pyruvate. HEK293T (ATCC® CRL-3216™), grown in DMEM 5% FBS, were used as packing cells for retroviral production to infect IECs for stable transformation.

Overexpressing plasmids were designed to carry a constitutive active form of the Akt isoforms. The plasmids were provided by adddgene; pBABE puroL (#1764) (Morgenstern y Land, s. f.), pBABE puroL Myr HA Akt1 (#15294) (Boehm et al. 2007), pBABE puroL Myr HA Akt2 (#9018)(Unpublished). For the lentiviral expression and stable transduction, we used pUMVC (#8449) and pCMV-VSV-G (#8454)(Stewart et al. 2003) addgene plasmids, into packing cells HEK293T. For transfection into the cells, we used Metafectene (Biontex) in a 1:3 (DNA: metafectene) ratio. Transfected cells were selected using Puromycin (Gibco) until growth was optimal, but it was removed during experimental assays.

### AKT INHIBITORS

Akt1 inhibitor, BML-257 (BML) Santa Cruz Biotechnology, CAS 32387-96-5 and Akt2 inhibitor, CCT-128930 (CCT) Santa Cruz Biotechnology, CAS 885499-61-6 were prepared on a stock solution of 10 mM on DMSO. The selected inhibitors were diluted to the indicated concentrations following the instructions provided by the suppliers. BML-257, further referenced as BML, is an Akt1 specific inhibitor which blocks the migration of Akt1 to the membrane to by activated. On the other hand, CCT128930, an Akt2 specific inhibitor, further referenced as CCT or CCT128, block the kinase activity of these isoform, so it blocks its downstream activity but not the capacity to be activated by mTOR, PI3K, PDK1…but does not decrease the activation phosphorylation on Ser474.

### WESTERN-BLOT (WB)

Protein samples were fractionated on SDS-PAGE, using Precision Plus Protein™ Dual Color Standards (BioRad) and nitrocellulose membranes (BioRad) and blocked in 3% BSA in TBS-T, and incubated with the primary antibodies at a 1:1000 dilution (pAkt1, pAkt2, No-p-βcat Ser45, p-βCatSer675 (CST), ZO1, Claudin4, Claudin2 Occludin, p-βCat Ser 33/37 (Invitrogen/Thermo Fisher), Claudin1, HSP90 (SantaCruz)) and secondary antibodies (Anti-Rabbit or Anti-Mouse, CST, 1:5000 and 1:2000) and revealed using Clarity Western ECL (BioRad). Image J was used to quantify the intensity of western blot bands. Each band intensity was corrected with a blank measure and further normalized to loading control.

### IMMUNOFLUORESCENCE

Cells were cultured on a slip. After the treatment, cells were fixed with 4% PFA for 30 minutes, permeabilized with TRITON X-100 0,1% on TBS-T and blocked in BSA 1% on TBS-T. Slips were incubated with the primary antibody, ZO1 (Thermo Fisher), Claudin2 (Thermo Fisher), LGR5 (AbCAM) or phospho-β-catenin (Ser33, Ser37) (inactive) (Thermo Fisher) in 1%BSA solution on a wet-chamber and with protection from light at 4°C. The next day, coverslips were incubated with mouse or rabbit Dylight 594 (Vector Laboratories) secondary antibodies and cytoskeleton markers, Phalloidin Alexa 647 (Thermo Fisher) and further incubated with DAPI (Merck, 1:7500). Finally, coverslips were mounted with Golden Prolong (Thermo Fisher) upside-down. Images were taken on confocal laser scanning microscope LSM900 (Zeiss) on vertical microscope Axio Imager 2 (Zeiss)

### MTT ASSAY

Confluent cell monolayers growing in medium without phenol red were treated with 250 ug/mL MTT (Sigma) in 96-well plates. After shaking slightly, they were incubated at 37°C and 5% CO2 for 4 hours until purple crystals were observed. Then, 90 μL of DMSO and then 60 μL of SDS 30% solution was added to each well. The plate was shaked, protected from light, for 10-70 min and then, optical density at 550 nm was measured on CLARIOstar OPTIMA (BMG LabTech)

### PERMEABILITY ASSAYS

For permeability assays (Liu et al. 2017; Zorraquín-Peña et al. 2021; Zufferey et al. 2009), cells were seeded on transwell plates (Corning 3413). After reaching confluence, they were treated for the indicated times and afterwards, 1 or 10 g/mL of FITC-Dextran 70 KDa (Sigma) was loaded on the upper chamber; 100 uL ofmedium from the bottom chamber were collected at either 1h or 4h after Dextran addiction and fluorescence was quantified (Emission; 490, Absorption; 520) on black 96 multiwell plates (Nunc) on CLARIOstar OPTIMA (BMG LabTech) Epithelial barrier resistance was monitored by transepithelial electrical resistance (TEER) assays; Briefly, cells were seeded on ECIS culture ware 8W10E-PET plates (Ibidi) and connected on ECIS Zo (Applied BioPhysics). After reaching confluence, the cells were treated and analyzed with the software provided by the supplier. (Meng y Roy 2016; Y. Gao et al. 2017)

### RT-PCR and RT-q-PCR

RNA was extracted using the commercial Nzyol (Nzytech) or TRIzol (Invitrogen) reagent and following the instructions provided. Reagents for RT and qPCR were purchased from Applied Biosystems and Promega, respectively and manufacturer instructions were followed. Human primers sequences (5’-3’); GAPDH Forward:<colcnt=6>

GCACAAGAGGAAGAGAGAG, GAPDH Reverse: AGGGGAGATTCAGTGTGGT, C-MYC Forward:

GCTGCTTAGACGCTGGATTT, C-MYC Reverse: TAACGTTGAGGGGCATCG, LGR5 Forward:

TCTGATCAGCCAGCCATC, LGR5 Reverse: CCCTTCATTCAGTGCAGTGT, Axin2 Forward:

TCTGTGGGGAAGAAATTCCATA and Axin2 Reverse: CAAACTCATCGCTTGCTTTTT. Mouse primers sequences (5’-3’); 18s Forward: GCAATTATTCCCCATGAACG, 18s Reverse: GGGACTTAATCAACGCAAGC, LGR5 Forward: TCCTGTCCATTTTTGCTTCC, LGR5 Reverse: TGACAGAAAAACCTCGTTCCA, Arg1 Forward: AGAGATTATCGGAGCGCCTT, Arg1 Reverse: TTTTTCCAGCAGACCAGCTT, NOS2 Forward: TGAAGAAAACCCCTTGTGCT, NOS2 Reverse: TTCTGTGCTGTCCCAGTGAG, CCL2 Forward: TTGACCCGTAAATCTGAAGCTAAT, CCL2 Reverse: TCACAGTCCGAGTCACACTAGTTCAC, IL1B Forward: TGTGAAATGCCACCTTTTGA and IL1B Reverse: GGTCAAAGGTTTGGAAGCAG.

### ORGANOIDS CULTURE

Mouse organoids were generate as described (Koledova 2017; Sato, van Es, et al. 2011; Sato et al. 2009, 5; Sato, Stange, et al. 2011). Briefly, male C57BL/6 mice 8–12-week-old mice were sacrificed by CO2 inhalation. To establish normal mouse colon organoids, crypts were isolated from colon tissue obtained after sacrifice. Briefly, the mouse colon was cut into small pieces, washed in PBS and incubated with antibiotic (Primocin) for 30 minutes. The tissue was then incubated for 5 min with 8 mM EDTA at room temperature and 30 min at 4 °C with slow rotation. The small pieces were then washed in PBS to remove EDTA. Crypts were isolated after shaking and collected after passing the solution through a 70 µm mesh filter. The crypt solution was centrifuged at 260 g for 5 min at 4 °C. Finally, organoids were cultured with Growth Factor Reduced (GFR) Basement Membrane Matrix (Matrigel), Phenol Red-free, LDEV-free (Corning) and grown with IntestiCult™ Organoid Growth Medium (Mouse) (Stem Cell Technologies). Organoids embedded in Matrigel were cultured on multi well chambers After the growth plus/minus treatments the chambers were washed and TRIzol (Invitrogen) added to each well. RNA and was extracted following the instructions provided by the manufacturer. Protein was extracted, using extraction buffer RIPA (50Mm TRIS HCl pH 7,4, 150Mm NaCl, 1% TRITON, 0,5% Deoxycholate, SDS 0,1%) with protease and phosphatase inhibitors (Roche) and 1 % NP-40 and 5 mM of EDTA and further sonicated on water on 15 cycles of 30 second ON and 30 second OFF. Then, protein concentration was measure using the BCA method, following the protocol provided by the supplier (Thermo Fisher)

For immunofluorescence (IF), organoids were cultured on micro-Slide 8-well IbiTreat (IBIDI) within 20 to 30 uL of Matrigel (Corning) containing the organoids. Structures were fixed with 2% PFA for 10 minutes, and permeabilizated and blocked in 1% DMSO, 1% BSA, 0,3 % TRITON X-100 on TBS-T. The next morning, organoids were incubated with mouse or rabbit Dylight 594 (Vector Laboratories) secondary antibodies and cytoskeleton markers, Phalloidin Alexa 647 (Thermo Fisher) and further incubated with DAPI (Merck, 1:7500). Images were taken on confocal laser scanning microscope LSM800 (Zeiss) on inverted microscope Axio Observer (Zeiss).

### MOUSE DSS MODEL

7-week-old C57BL/6 mice male mice were originally obtained from Charles Rivers and bred and maintained in the CBMSO facilities. Colitis was induced with DSS (Chassaing et al. 2014) added to drinking water at a concentration of 3% and given at libitum for a week. Weight was monitored every day. Akt inhibitors CCT128930 (Cayman) and BML257 (Enzo) were administered every day via intraperitoneal injection. Every compound was diluted following the manufactureŕs instructions to a concentration of 10 mg/mL each. Each animal received a final dose of 1mg daily(Yap et al. 2011; O. Chen et al. 2017). Disease severity was analyzed following previous guidelines (Sann et al. 2013)

For colonoscopy, mice were anesthetized with Isoflurane (concentration) and laid on supine position with isoflurane intake. Colonoscopies were carried out in an endoscope (Karl Storz) as described (Becker, Fantini, y Neurath 2006; Becker 2005; Kodani et al. 2013).

All mice were kept under Specific Pathogens Free (SPF) conditions in the Animal Facilities at the CBMSO following national and European guidelines (PROEX004/17, PROEX240/19). Every procedure was approved by the ethics committee at the CBMSO. Animals were euthanized by CO2 inhalation at the end of each experiment. In case of loss of quality of life by a direct or indirect consequence of the experiment, mice were sacrificed.

### TISSUE IMMUNOHISTOCHEMISTRY AND TUNNEL ASSAY

Colon rolled paraffin slides were incubated for antigen unmasking with Antigen Retrieval (DAKO), with ABC kit (VectorLabs). Slides were incubated with anti Akt1 and Akt2, (Sigma, 1:250), ZO1 and Claudin4 (Invitrogen/ThermoFisher, 1:250), EPCAM (CST, 1:5000), Claudin1 (SantaCruz, 1:200) and the bound secondary antibody – peridoxidase (DAKO) was detected with Liquid DAB + substrate chromogen system (DAKO) following the instructions provided by the manufacturer. Slides were counterstained with Hematoxylin solution (Sigma) for 10 seconds and mounted with DPX new (MERCK). Slides were also stained for conventional histological studies with Hematoxilin/Eosin and Gomori Trichrome staining. Images were taken on a vertical microscope Axioskop2 plus (Zeiss) with a DMC6200 camera (Leica).

For TUNEL assays, paraffin slides were incubated with the Click-iT Plus TUNEL assay kit (Invitrogen) following the instructions given by the manufacturer, with the exception that DAPI (Merck, 1:5000) was used instead of Hoechst. Images were taken on a confocal laser scanning LSM900 (Zeiss) on a vertical microscope Axio Imager 2 (Zeiss).

### IL10 ELISA

Serum was obtained from mice after 1h incubation of blood samples at 37°C and at least 2h at 4°C followed by a centrifugation step at 1500 rpm for 5 minutes. ELISA was performed following the instructions provided by the manufacturer (Mouse IL-10 DuoSet ELISA; R&D systems) and read out on CLARIOstar OPTIMA (BMG LabTech).

### STATISTICAL ANALYSIS

The statistical evaluation was done using GraphPad Prism 8-9; Statistical significance determined without correction for multiple comparisons, with alpha=0.05%. Each gene was analyzed individually, without assuming a consistent SD. ANOVA test were performed. Results are expressed as mean ± SEM. The results were considered statistically significant when the value of the p-value was equal to or less than 0.05, represented as*p<0,05, **p<0,01, *** p<0,001, ****p<0,0001.

## RESULTS

### Akt1 and Akt2 differentially affect epithelial barrier permeability and tight junction protein expression

To investigate how Akt isoforms affect the IEC, we first stably overexpressed either Akt1 (Akt1+) or Akt2 (Akt2+) on several IEC lines. We used at least 3 IEC for each experiment to avoid obtaining particular results with a single IEC. The results indicate that Akt1 overexpression led to an increment on membrane cell resistance in TEER assays HT-29 (Fig 1a), HCT116 (Fig. S1a) and SW480 (data not shown) cell lines. Akt1+ cells also showed a decrease on the cell monolayer permeability to FITC (Fig. 1b). In contrast, overexpression of Akt2 had no major effect. We addressed whether those changes in barrier function correlated with alterations in TJP expression in the epithelial cells. A variable upregulation on the levels of barrier-forming TJP, such as Claudin1 and ZO1 was observed by western blot in different IEC lines HT-29, HCT116, and SW480 Akt1+ (Fig. 1c, d and Fig. S1c, d). In contrast, Akt2+ cells have the opposite phenotype with a tendency to decrease on those TJP levels. By IF, we further confirmed that ZO1 expression is not only increased, but more importantly, it also seems to be located on the membrane of Akt1+ cells, suggesting changes on permeability (Fig. 1e and Fig. S1e). In contrast, anti-barrier Claudin 2 is more expressed on Akt2+ cells (Fig. 1e and S1e). Taken together, these results suggest that Akt1 is important on the maintenance and functionality of the intestinal epithelial barrier.

**Figure 1.**
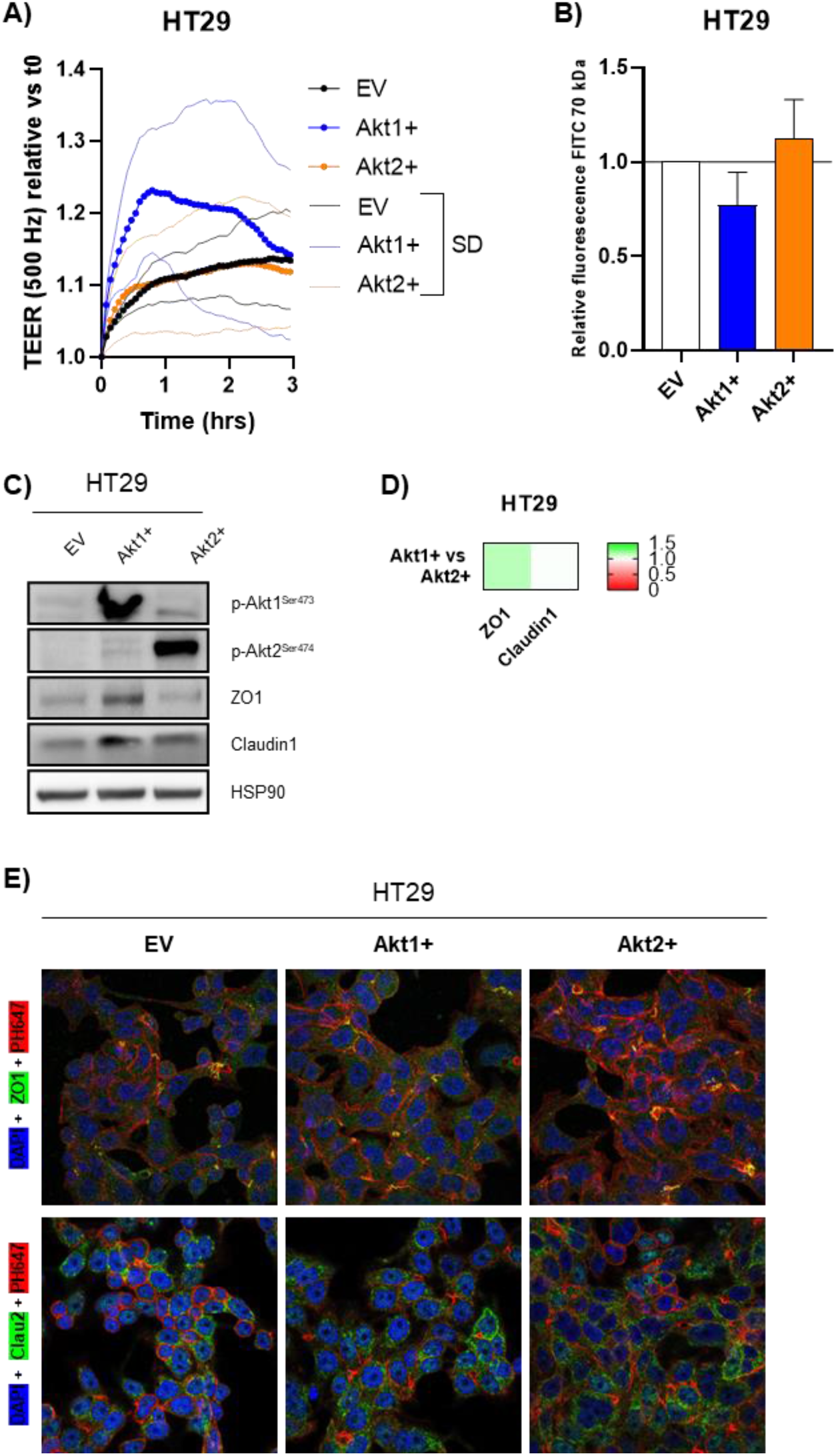
Akt1 and Akt2 overexpression differentially affects epithelial barrier resistance and TJP expression in the HT29 IEC line. (A) TEER assays were performed to determine the barrier resistances of the overexpressing cells. Cells overexpressing Akt1 and Akt2 are labeled as Akt1+ or Akt2+. (B) In parallel, permeability of the epithelial barrier was assayed with FITC-Dextran 70KDa. (C) Western blot analysis of ZO1, and Claudin1 in the overexpressing cell lines and (D) quantification of 3 WBs; each band density was corrected by HSP90 expression. Then Akt1+/Akt2+ ratio of expression was represented. (E) Expression and localization of ZO1 by IF; Nucleus (Blue, DAPI), Actin peripheral cytoskeleton (Red, Phalloidin) and ZO1 (Green).

Next, we performed the opposite approach by inhibiting the different isoforms with the use of specific pharmacological inhibitors (BML-257 for Akt1 and CCT-128930 for Akt2). TEER and permeability assays performed on several IECs demonstrated that treatment with the Akt2 inhibitor increased the epithelial barrier resistance (Fig. 2a and Fig. S2a) and decreased FITC membrane permeability (Fig. 2b and Fig. S2b). In contrast, treatment with the Akt1 inhibitor strongly decreased the membrane resistance in most of IEC lines and increased the FITC permeability, providing a reduced barrier function. An increase on the levels of barrier forming TJP such as Occludin and ZO1 and a decrease on the pro-leaking proteins Claudin2, was also observed in cells treated with the Akt2 inhibitor CCT within just 3 minutes (Fig. 2c, d and Fig. S2c, d). Moreover, CCT-treatment resulted in enhanced degradation of anti-barrier Claudin2. Interestingly, this profile was similar to the one observed in Akt1+ cells. This was further supported by the fact that Akt2-inhibited cells with CCT present an increase on active Akt1 levels. On the other hand, cells treated with BML, an inhibitor of Akt1 translocation that prevent this kinase to become activated, presented less significant changes but there was a slight decrease on the previously mentioned TJP (with the same trend as was observed in Akt2 overexpressing cells). Moreover, in IF assays, Akt2 inhibitor-treated cells showed an increased expression and colocalization of ZO1 with the actin peripheral cytoskeleton (Fig. 2e and Fig. S2e), suggesting ZO1 is more abundant on the cell membrane, where it can act in the tight junctions. In contrast, treatment with the Akt1 inhibitor BML induced an increase of Claudin2, a leaking and pore-forming protein, on the membrane (Fig. S2e).

**Figure 2.**
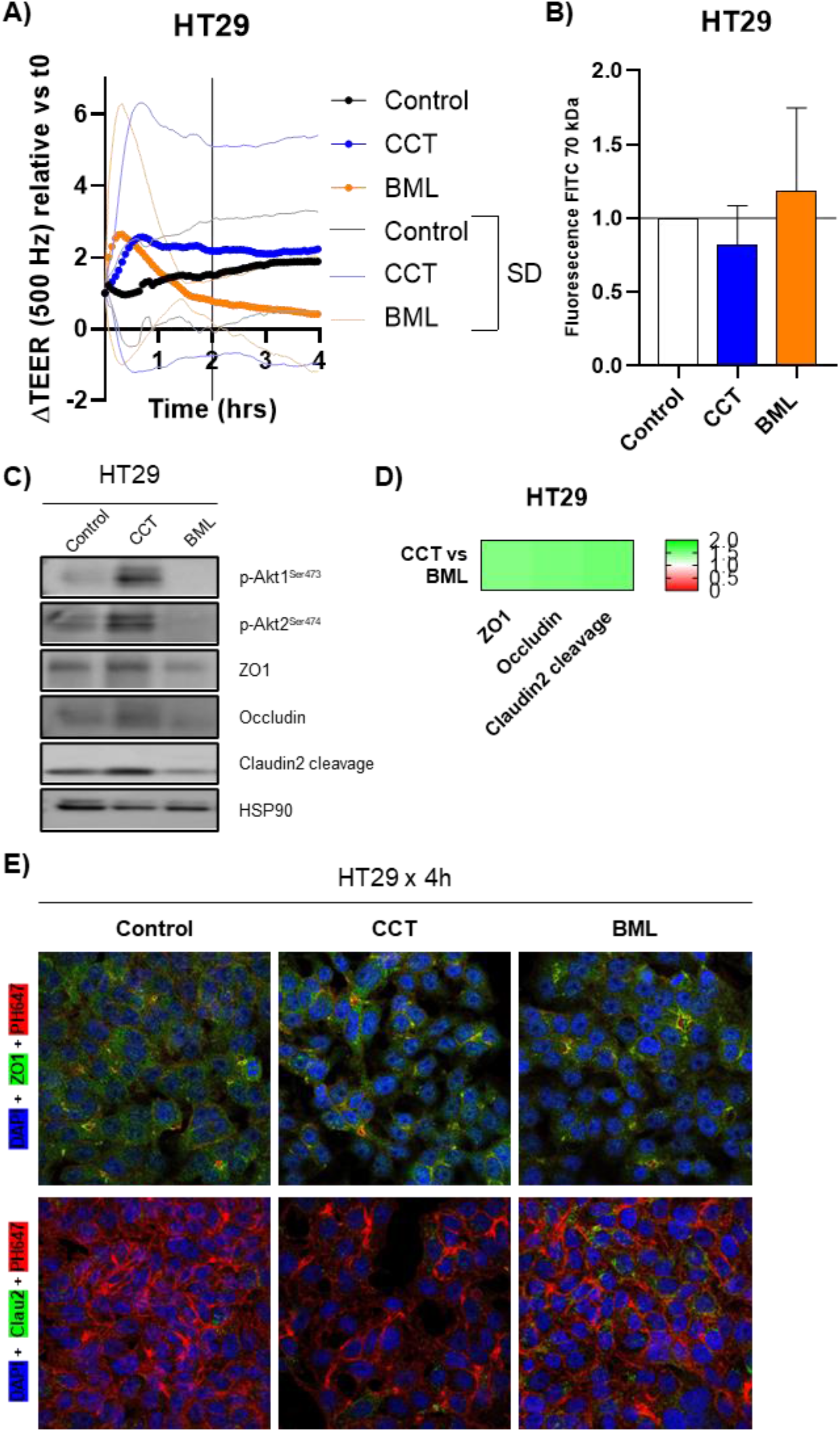
Pharmacological inhibition of Akt2 increases intestinal epithelial barrier function. HT29 IECs were treated with either CCT128 (Akt2 inhibitor), or BM257 (Akt1 inhibitor), or ATP-competitive (negative control). (A) TEER assay was performed to determine the barrier membrane resistance of cells. (B) Permeability assay with FITC-Dextran 70 KDa 1h post FITC incubation. HT29 confluent cells were treated with both inhibitors for 30 minutes and tight junctions’ proteins and Akt phosphorylation were checked by (C) western blot and (D) quantification of 3 WBs; each band density was corrected by HSP90 expression. Then CCT/BML ratio of expression was represented. (E) IF staining of TJP’s in HT29 cells after treatment with CCT128 and BM257, nucleus (Blue, DAPI), Actin peripheral cytoskeleton (Red, Phalloidin) and ZO1, top, or Claudin2, bottom, (Green).

### Akt isoforms control differently β-Catenin phosphorylation and activation of WNT pathway

Another major regulator of intestinal barrier homeostasis is the cell renewal on the crypts, process in which the β-Catenin/WNT pathway is involved. Thus, we further investigated how Akt1 and Akt2 affect the survival and regeneration of the IECs. Both Akt1 overexpression and Akt2 inhibition resulted in an increase on active β-Catenin levels (Phospho Ser675 and Non-phospho Ser45) detected by WB (Fig. 3a,b and Fig. S3a,b). These two modifications are considered as markers of translocation of this protein to the nucleus, where it can act as a transcriptional factor, promoting survival and regeneration of the epithelia. On the contrary, Akt2+ cells and Akt1 inhibited cells present an increase on inactive β-Catenin (Phospho Ser33/37 and Phospho Thr41) (Fig. 3a,b and Fig. S3a,b). Phosphorylation in these sites promotes β-Catenin degradation, resulting in a decrease in cell survival. In agreement with that, IF staining showed that HCT116 Akt1+ cells had much lower levels of inactive βCat (Fig. 3c) and HT29 Akt1+ cells had higher levels of LGR5 (Fig. S3c), a main proliferation marker. Analyses of the membrane to nucleus ratio show a significant redistribution of inactive β-Catenin, being relatively higher in the membranes of Akt1+ cells but lower.in Akt2+ cells that in EV control cells (Fig. 3d). The opposite relative cellular distribution was observed with LGR5 (Fig. S3d).

**Figure 3.**
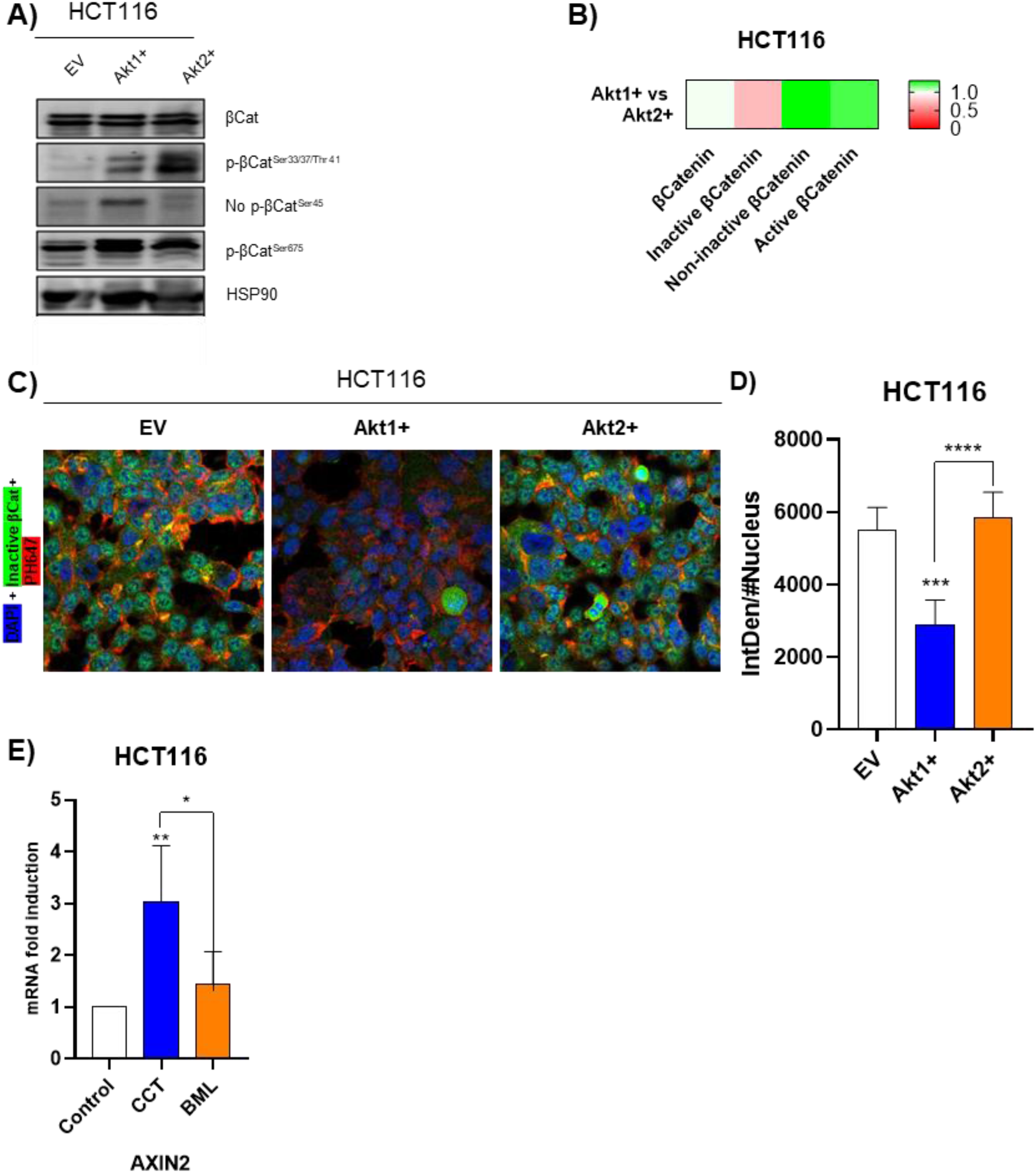
Akt isoforms control β-Catenin phosphorylation and WNT pathway. The HCT116 IEC line was stably transfected to overexpress Akt1 and Akt2 (A) Analysis of the state of activation of β-Catenin in HCT116 Akt1+and Akt2+ cells by western blot (active: p-βCat Ser675 and No p-βCat Ser45; marked to degradation: p-βCatSer33/37/Thr41) (B) quantification of 3 WBs; each band density was corrected by HSP90 expression. Then Akt1+/Akt2+ ratio of expression was represented. (C) IF staining of p-βCatSer33/37/Thr41 in HCT116 Akt1+ and Akt2+ cells, marking nucleus (Blue, DAPI), Actin peripheral cytoskeleton (Red, Phalloidin) and inactive β-Catenin (Green) and (D) IF quantification of inactive βCatenin intensity corrected by number of cells (Nucleus). (E) qPCR for *Axin2* in HCT116 IECs after treatment with Akt inhibitors for 24h.

Besides, we checked the activity of β-Catenin as a transcription factor through monitoring mRNA levels of *Axin2* and *cMyc*, which are targets of the WNT pathway, being its induction related to survival promotion. Inhibition of Akt2 strongly increased *Axin2* and *cMyc* mRNA levels after 24h of administration of CCT, whereas BML treatment, an inhibitor of Akt1, had no significant effect (Fig. 3e and Fig. S3e). We further checked the viability of HT29 cells in MTT assays after 24h of inhibition and observed a slight but significant decrease on cell survival on those cells with the Akt1 inhibitor BML and a slight increase on Akt2-inhibited cells (Fig. S3f).

### Akt inhibitors modulate organoidś structure and inflammatory status

To further determine the role of Akt1 and Akt2 on intestinal inflammation, we checked the effect of Akt inhibitors on a more physiological system, mouse colon intestinal organoid. In non-treated colon normal organoids, the cells are oriented, and show a cortical actin cytoskeleton in the inner part of the organoid that corresponds to the apical zone. We observed that the inhibition of Akt1 with BML led to a disruption of the actin cytoskeleton, resulting in a depolarization, a reduction on the number of polarized “oriented” cells giving rise to compact columnar” organoids without lumen. Those BML-treated organoids present stratified layers of cells (Fig. 4a). Organoids with a decrease in “oriented” cells and “columnar” organoids, are indicators of diseased or “IBD-derived” organoids (d’Aldebert et al. 2020). In contrast, Akt2-inhibited organoids remain mostly unchanged, although an increase on organization was observed (Fig 4a,b).

**Figure 4.**
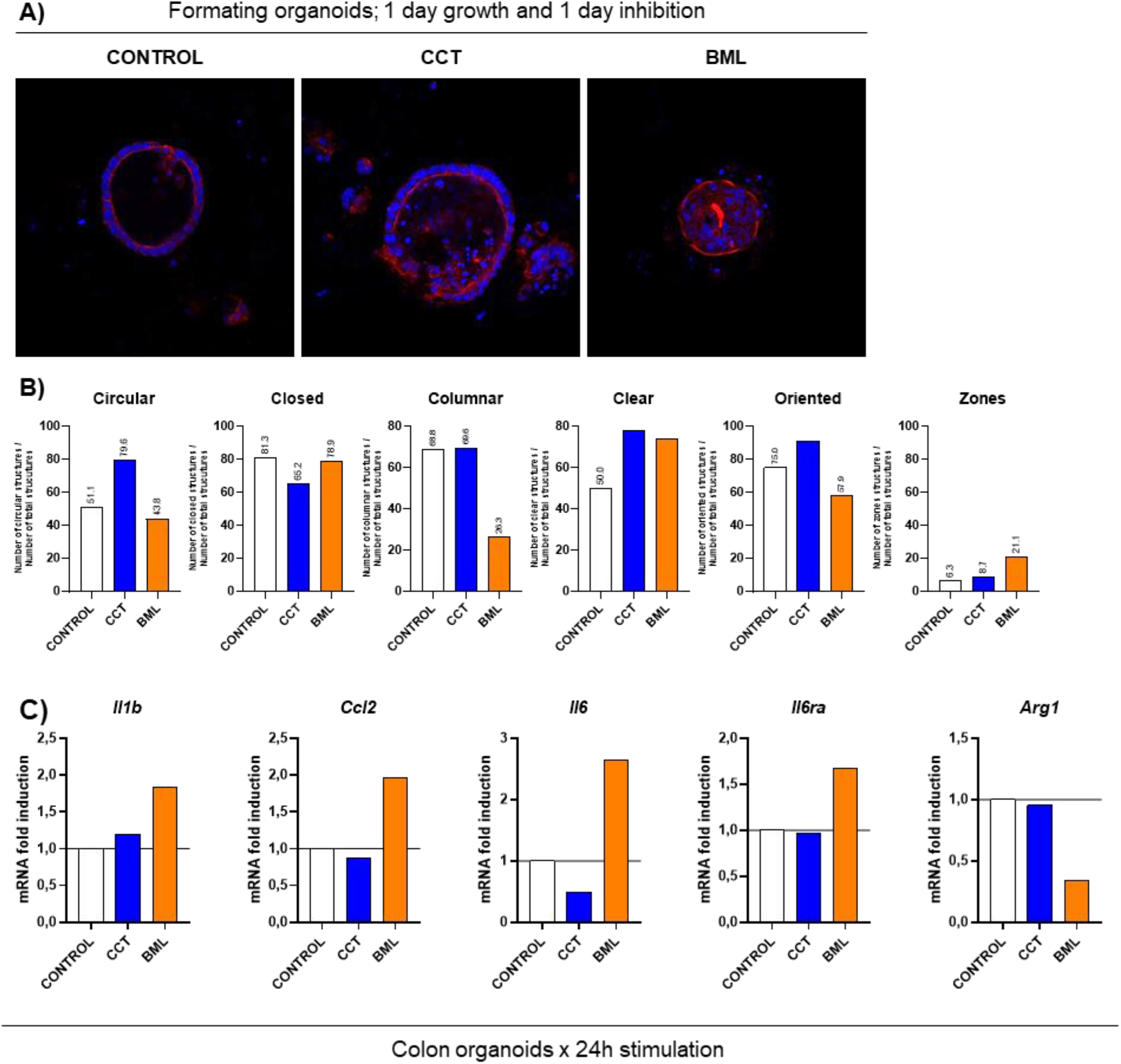
Akt inhibition modulates organoidś structure and inflammatory environment. Mouse organoids were cultured for 24h after colon crypts isolation and then, cultures were treated with AKT1 (BML) and AKT2 (CCT) inhibitors for 24h; F analysis of organoidś structure (A) representative image marking nucleus (Blue, DAPI) and Actin peripheral cytoskeleton (Red, Phalloidin) and (B) quantification of structures. (C) qPCR analysis of pro and anti-inflammatory mediators’ mRNA levels on the organoids in response to treatment with the Akt inhibitors.

We also analyzed gene expression of inflammatory markers by qPCR, showing that Akt1-inhibited organoids present an increase on the mRNA levels of the pro-inflammatory mediators *Ccl2, Il6/IL6ra* and *Il1b,* and a decrease on the anti-inflammatory marker *Arg1* (Fig. 4c). On the other hand, on Akt2-inhibited organoids, we could only observe a decrease on *Il6* mRNA levels, suggesting that Akt1 seems to be more important than Akt2 on the regulation of inflammation in colon, in absence of infiltrating cells.

### Akt1 and Akt2 inhibitors play opposite roles on IBD development in mice

Our previous results indicate an important and differential effect of AKT isoforms on the intestinal epithelial cell barrier and organization, which are markers of IBD development. Thus, we wanted to determine if treatment with Akt inhibitors would affect the development of ulcerative colitis in mice. Colitis was induced with DSS and mice were organized into four groups: control healthy mice, control DSS mice, and DSS mice treated with either Akt1 inhibitor or Akt2 inhibitor. The progression of the disease was monitored through a disease score (DAI) (Fig 5a) and also by a less subjective novel non-invasive technique, colonoscopy, that allows continuous real time monitoring and reduces the numbers of sacrificed animals needed (Fig. 5b and 5c). Control DSS treated animals start to develop significant signs of disease after 4 days as observed by both techniques (Fig 5a and not shown respectively). Interestingly, BML-treated mice presented a significant acceleration of the progression of disease in the first 5 days with abundant diarrhea (Fig. 5a) and increasing ulcers and thickness of the intestinal wall (Fig 5c). In contrast, and even more interestingly, Akt2-inhibitor treated mice showed a reduction in the development of the disease and a much less ulcerated colon, which was evident in the colonoscopy analysis (Fig. 5b,c). In agreement with the disease scores, histological analysis of colon samples showed a strong damage in response to DSS treatment that was much worsened in DSS mice treated with the Akt1 inhibitor; these animals present less crypts, and more inflammation and ulceration than control diseased DSS mice (Fig. 5d). In contrast, the inhibition of Akt2 resulted in a strong improvement in the histological characteristics of the intestinal wall, with more well-formed crypts and less inflammation on the colon tissue. Gomori Trichrome staining (Fig. 5e) allows differentiating connective tissue (collagen in blue) from muscle fibers, crypts and epithelial tissue (in red). Large infiltrations of inflammatory cells on the mucosa and submucosa and enlargement of the submucosa were present, which are the main characteristics of IBD due to exacerbated inflammation. In mice treated with the inhibitor of Akt1, we observed no crypt’s structure and a thicker submucosa even than in DSS control mice. In contrast, treatment with the Akt2 inhibitor, CCT, resulted in basically no enlargement of the submucosa and the tissue structure was more similar to control healthy mice than to control DSS mice (Figure 5d and e). Altogether, those results indicate that the inhibition of Akt1 enhances the development of the disease whereas the inhibition of Akt2 ameliorates it. Furthermore, tunnel assays were performed to evaluate in situ apoptosis and cell death due to damage. Apoptosis was not as extensive on the tissue of animals treated with the Akt2 inhibitor compared to diseased control mice. In animals treated with the Akt1 inhibitor, the incidence of apoptosis was even higher (Fig. 5f).

**Figure 5.**
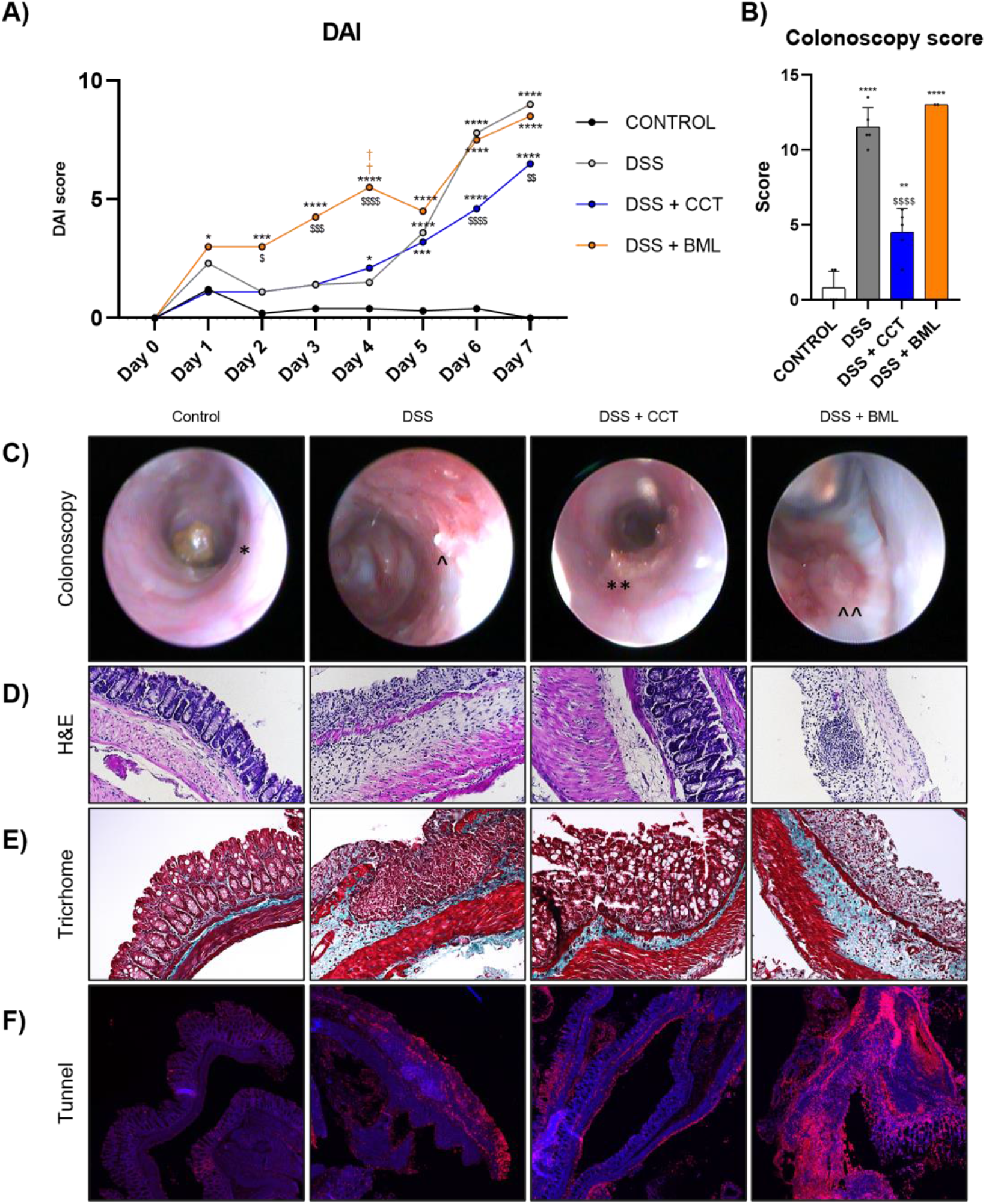
Akt1 and Akt2 inhibitors play opposite roles on IBD development in mice. Mice were treated with 3% DSS on drinking water at libitum (DSS, disease control mice) and treated with either Akt2 inhibitor CCT128 (DSS + CCT) or Akt1 inhibitor BML257 (DSS + BML), for a week. CONTROL stands for mice given control water and vehicle treatment. (A) DAI score for the four groups of mice. (B) In vivo colonoscopy score showing the progression of the disease at final day and (C) correlated images, marking mean features; *=formatted stool, **= unformatted stool, ^= Redness ulcer and ^^= Bleeding ulcer. (D) H&E and (E) Gomori trichrome staining of the tissue postmortem to evaluate the damage on the colon tissue. (F) Tunnel staining of the tissue to check apoptosis in situ. * stands for p-value vs Control and $ vs DSS.

Besides, immunocytochemical staining for EpCAM, a main intestinal epithelial marker, showed a dramatic reduction on the number of crypts and epithelial cells after DSS treatment. This was reversed to almost normal non-diseased levels by treatment with the Akt2 inhibitor (Fig. 6a). Immunocytochemical staining for ZO1 (Fig. 6b), Claudin1 (Fig. S6c) and Claudin4 (Fig S6d) indicates that treatment of DSS animals with the Akt2 inhibitor not only helps maintain the integrity of the intestinal crypts and tissue architecture, but also ensures that the epithelial cells in the crypts were correctly bound to each other as detected by the presence of those barrier-forming proteins all along the crypts. Altogether, these results indicate that the organization, structure of the colon and TJP expression are close to normal in DDS diseased mice treated with the Akt2 inhibitor.

**Figure 6.**
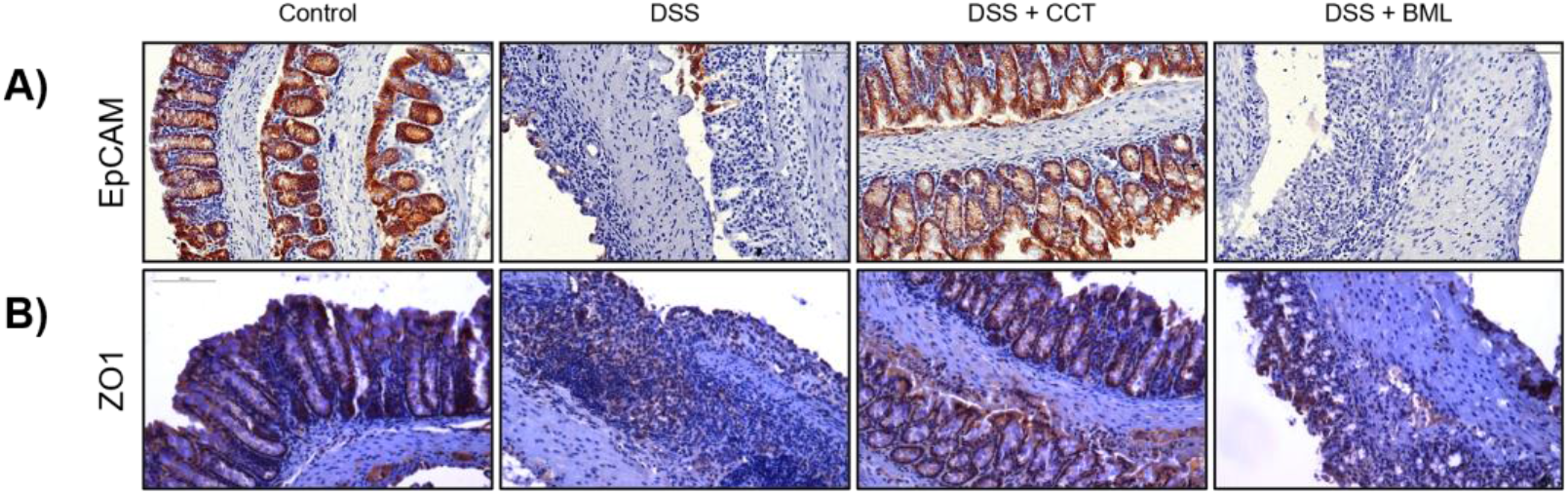
Treatment with Akt1 or Akt2 inhibitors differentially affects epithelial junctions upon IBD induction. Mice were treated with DSS 3% on drinking water ad libitum (DSS, disease control mice) and treated with either Akt2 inhibitor, CCT128 (DSS + CCT) or Akt1 inhibitor, BML-257 (DSS + BML), both at 1 mg/mouse, each day for a week. Control mice were given control water and vehicle treatment. Immunohistology staining of (A) EpCAM, to check the crypts structure, (B) ZO1, as pro-barrier tight junction proteins.

Interestingly, Akt1 immunostaining of the tissues demonstrated that Akt1 is present in the colon tissue of mice in the four experimental conditions, with a higher intensity of immunostaining and a higher number of positive cells in animals treated with the Akt2 inhibitor (Fig. S6a). However, Akt2 was almost absent on control mice colon, but it was expressed after induction of IBD by DSS, especially on the submucosa infiltrates (Fig. S6b). Interestingly, Akt2-inhibitor treated mice had a healthier colon and, consequently, Akt2 staining was almost absent.

### Akt1 inhibition creates a pro-inflammatory environment as opposed to Akt2 inhibition

IL10 serum levels, a main anti-inflammatory regulator of IBD development (Iyer y Cheng 2012, 10) decreased with the DSS treatment, as expected, and this decrease was also not significantly modified in animals treated with the Akt2 inhibitor, but it was even higher in animals treated with the Akt1 inhibitor (Fig. 7a). The balance of Arg1/Nos2 mRNA levels in the spleen can be taken as a surrogate marker of M2/ M1 ratio (Jablonski et al. 2015). As described before (Eichele y Kharbanda 2017; Dieleman et al. 1998), DSS resulted in a more proinflammatory ratio (Fig. 7b). Treatment with the Akt2 inhibitor resulted in a M2-like environment, even higher than healthy controls, whereas treatment with the Akt1 inhibitor resulted in an even more pro-inflammatory, M1-mediated environment. Analyzing several pro and anti-inflammatory mediators’ mRNA levels on colon tissue revealed that Akt1 inhibition increased the pro-inflammatory (*I1lb, Ccl2*) and decreased the anti-inflammatory (*Arg1*) mRNA levels, while Akt2 inhibition increased the anti-inflammatory environment as well as decreased pro-inflammatory mRNA levels (Fig. 7c). Lgr5 expression, a main epithelial stem cell marker, on colon tissue decreases with the IBD induction but decreases even more with the inhibition of Akt1 (Fig 7d), being those cells likely incapable of self-renewal and restock the crypts lost with DSS.

**Figure 7.**
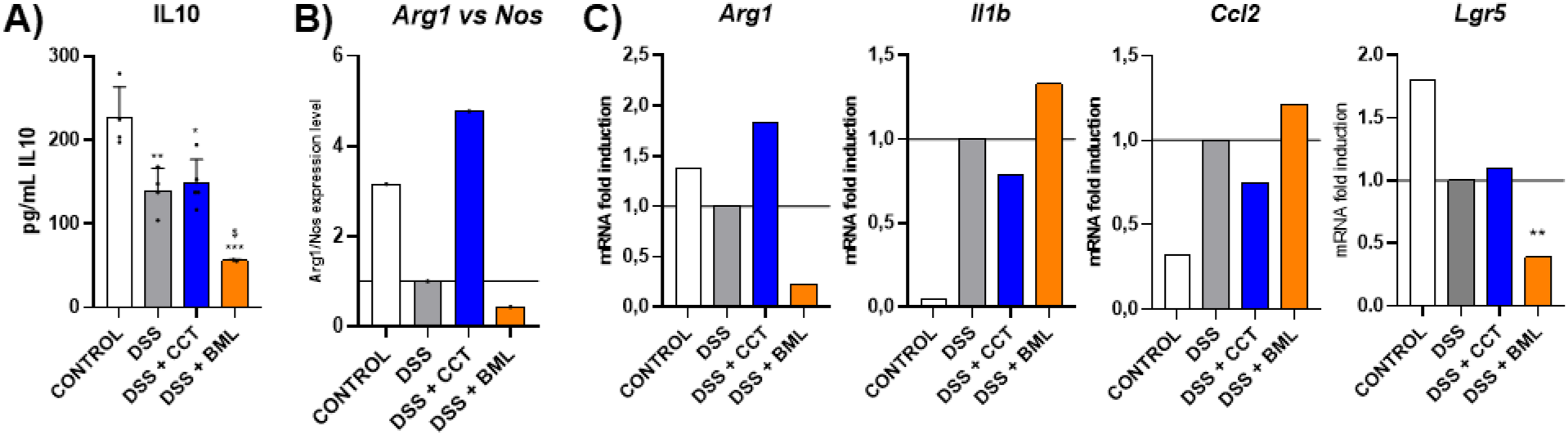
Akt2 inhibition creates an anti-inflammatory environment opposite to the inhibition of Akt1. (A) Serum levels of IL10 as determined by ELISA. (B) Ratio between M2 marker *Arg1* and *Nos2*, M1 marker, to determine M1 or M2 phenotype by qPCR on spleen tissue. (C) Immunological markers *Arg1, Il1b, Ccl2* and Lgr5 stem cell marker, mRNA expression levels by qPCR analysis of on colon tissue.

## DISCUSSION

Our results show that treatment with an Akt2 inhibitor prevents the development of IBD in mice and improves the integrity of the intestinal epithelial barrier, whereas treatment with an Akt1 inhibitor has the opposite effect. This is in agreement with previous studies showing that Akt2-/-mice were more resistant to DSS-induced colitis than wild-type mice, whereas Akt1-/-mice were more sensitive (Arranz et al. 2012). However, that study focused on the effect of the Akt isoforms only in the immune cells, mainly macrophages. Thus, Akt kinases were shown to differentially contribute to macrophage polarization, with *Akt1* ablation giving rise to an M1 phenotype and *Akt2* ablation resulting in an M2 phenotype. Moreover, reconstitution of irradiated WT mice with Akt2-/-bone marrow only partially reduced the severity of DSS-induced inflammatory bowel disease (Arranz et al. 2012), suggesting that the M1 or M2 phenotype is important in the development of IBD but it is not the only cause of the disease. On the other hand, another important player for IBD development is the integrity of the intestinal epithelium, where Akt1 and Akt2 can be expressed and thus may play a role in its physiopathology. In this regard, IBD can be caused by genetic mutations that predispose the dysfunction of the intestinal mucosal barrier and some of those genes can affect the development of IBD through signaling pathways that involve PI3K/Akt activation (Wei y Feng 2010).

We have shown here for the first time that the inhibition of Akt2 in vitro increases the intestinal barrier function, thus decreasing its permeability, and we have also confirmed an increased anti-inflammatory environment (or M2 phenotype) when Akt2 is inhibited. All those features contribute to protection against development of IBD. In contrast, we found that Akt1 inhibition increases permeability and decreases epithelial barrier function, as well as promotes inflammation (or M1 phenotype). In addition, we found that Akt1 inhibition may affect the renewal Lgr5+ IEC by turning off the β-Catenin/WNT pathway and so further increasing IBD development. In this regard, there are a few reports showing differential roles of Akt isoforms in various disorders including cardiovascular diseases or cancer (Yu, Littlewood, y Bennett 2015; Cohen 2013; Hinz y Jücker 2019).

The relevance of our study relies on the opposing roles of Akt1 and Akt2 on the intestinal epithelium. The intestinal barrier and its correct regulation and permeability is a key feature for the correct function of the intestine, and a disruption of this barrier leads to IBD development. We demonstrated here that Akt2 is implicated on the negative regulation of this process, since Akt2 inhibition led to an increase on barrier-forming TJPs and a decrease of pore-forming proteins such as Claudin2, resulting in a more tightened barrier and a decrease on epithelial permeability. In this regard, it has been shown that Akt affects the interaction of Connexin43 with ZO-1, allowing gap junctions to enlarge (Solan and Lampe 2014) and that Akt DN induces ZO-1 (Xiao et al. 2015). However, these studies did not point to which of the isoforms of Akt was involved in the regulation of the TJP. We have found that Akt1 is a positive regulator of intestinal permeability, since its inhibition led to a dysregulation of the barrier and its overexpression maintained the functionality of the intestinal epithelial barrier. Altogether, our results suggest that the active Akt1 isoform should be found in a higher proportion than the Akt2 isoform in order to control the tightness of the intestinal epithelial barrier and its function. Another important feature of our studies was to demonstrate the cross-regulation between both Akts. This was supported by the fact that pharmacological inhibition of Akt2 with CCT resulted in a huge increase on active Akt1 levels. This cross-regulation, may also help explain some paradoxical effects reported with this CCT inhibitor, as it has been described that CCT induces activation of downstream effectors of Akt in HEPG2 hepatoma cells (Wang et al. 2014). This negative crosstalk between Akt1 and Akt2 results in the similar effects observed either overexpressing AKT1 or inhibiting AKT2.

Another major player which affects the progression of IBD is the renewal and survival of IECs (Okumura y Takeda 2017; Okamoto y Watanabe 2016). A key player in this is the Wnt/β-catenin pathway (Moparthi y Koch 2019). We demonstrated that both overexpression of Akt1 and inhibition of active Akt2 increased the activation of the β-Catenin/WNT pathway. Thus, β-Catenin is phosphorylated on Ser552 and Ser675, activation residues that mark its translocation to the nucleus where it acts as a transcriptional factor, increasing the transcription of pro-survival factors such as c-Myc or Axin2, a major stem cell marker, and LGR5. In contrast, Akt1-inhibited or Akt2-overexpressing cells presented a decrease on cell survival correlated with phosphorylation on Ser33/37 and Thr41, which mark β-Catenin for degradation, resulting in a decrease of active βCatenin. In addition, mice treated with the Akt1 inhibitor also presented a decrease in the Lgr5 mRNA levels in colon tissue. Activation of the β-Catenin/WNT pathway induces Lgr5+ stem cells to promote epithelial reconstitution in experimental colitis ameliorating the disease(Luo et al. 2022). Besides, elimination of Lgr5+cells resulted in crypt loss (Metcalfe et al. 2014). Our results suggest that Akt1 plays an important role on cell survival and its absence leads to a severe. Lrg5 depletion in vivo and thus to the inability of cell renewal after IBD intestinal damage. Opposing effects of Akt1 and Akt2 on β-Catenin have been already described (S.. P. Gao et al. 2021; Mauro et al. 2009). In cancer, Akt1 has been shown to impair apoptosis (Hinz y Jücker 2019), which is consistent with our observed effects on cell survival. Thus, the ratio of active Akt1/Akt2 may be important for the activation of βCatenin/WNT pathway (S. P. Gao et al. 2021; Mauro et al. 2009).

On mouse organoids, we found a direct effect of Akt1 in intestinal epithelial organization. Inhibition of Akt1 led to a depolarization and disorganization of the structures and at the same time to an increase on pro-inflammatory mediator’s mRNA levels and a decrease on the anti-inflammatory ones. Altogether, these results suggest that Akt1 is the most relevant isoform in the regulation of the intestinal epithelial barrier organization.

Finally, we have shown in vivo, in a DSS-induced colitis model in mice, that Akt1 and Akt2 have a differential role on IBD development. Thus, Akt1 inhibition promotes the development of the disease, with colon samples showing more tissue damage, less intestinal crypts and more immune infiltration on the submucosa. Furthermore, Akt1 inhibition led to decreased IL10 on serum, and spleen macrophages showed a pro-inflammatory M1 phenotype, as well as to an elevation on mRNA levels of inflammatory markers and a decrease on anti-inflammatory molecules on colon tissue. In contrast, Akt2 inhibition had an opposite role in most of the experiments performed. Thus, mice treated with the Akt2 inhibitor showed much lower intestinal damage, and the histological appearance of colon tissue was much closer to the one observed in healthy mice. Moreover, the negative implication of Akt2 in DSS-induced colitis is supported by the fact that Akt2 expression is strongly induced in the damaged colon and in the infiltrated cells after DSS treatment.

In summary, our results reported here point out a balance between Akt1 and Akt2 activation regulating IBD development through several mechanisms, being Akt1 and Akt2 differentially involved in some of the mechanisms in IBD, as intestinal permeability, inflammation, or IEC renewal. When Akt1/Akt2 is diminished, the intestinal epithelial barrier is disrupted by the dysregulation of the TJPs, leading to an acute phase of inflammation. Moreover, the intestinal epithelium is unable to renew itself properly in the absence of active Akt1, making the damage more and more severe, and creating a pro-inflammatory niche that derives on chronic inflammation and severe IBD. In addition, AKT1 may be involved in a better self-renewal of the damaged crypts IBD. Furthermore, the inhibition of Akt2 may be considered as a treatment or adjuvant treatment for IBD. In this regard, the PI3K/AKT/PTEN pathway has been already suggested as a target for Crohn’s disease therapy (Tokuhira et al. 2015). Besides, our results on the different actions of the two isoforms underline the importance of clinical treatments targeting specific Akt isoforms, as it has been suggested for other human diseases, mostly cancer (Heron-Milhavet et al. 2011). Finally, given the importance of Akt2 and its enhanced expression in IBD, as described here, Akt2 may be considered as a new marker of the progression of the disease.

---

**Supplementary figure 1.**
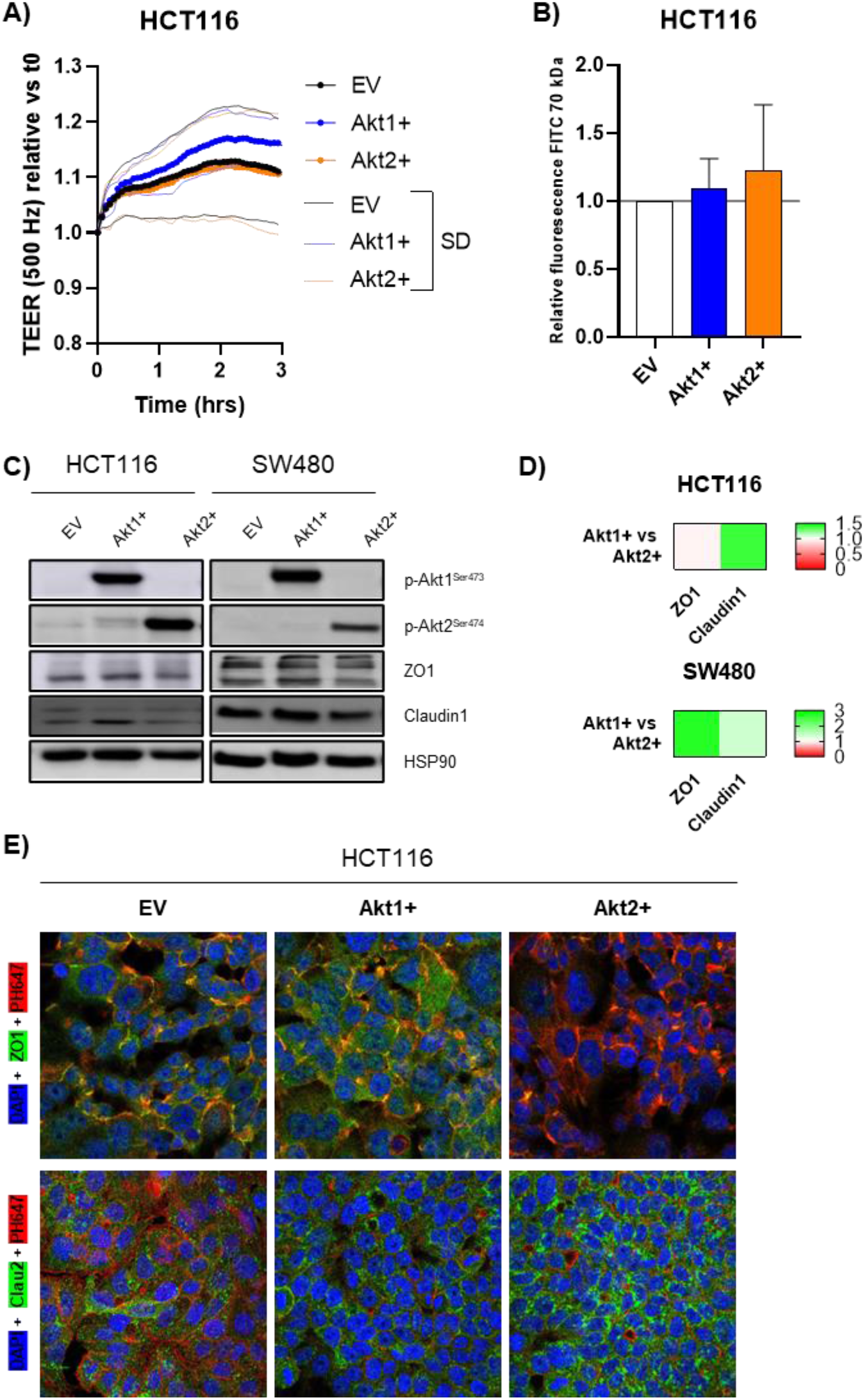
Akt1 and Akt2 overexpression differentially affect epithelial barrier resistance and TJP expression. HCT116 and SW480 cells overexpressing Akt1 and Akt2 are labeled as Akt1+ or Akt2+. (A) TEER assays were performed to determine the barrier resistances in HCT116 overexpressing cells. (B) In parallel, permeability of the epithelial barrier was assayed with FITC-Dextran 70KDa in HCT116 overexpressing cells. (C) Western blot analysis of ZO1 and Claudin1 on HCT116 and SW480 cells overexpressing Akt1 (Akt1+) or Akt2 (Akt2+), and (D) quantification of 3 WBs; each band density was corrected by HSP90 expression. Then Akt1+/Akt2+ ratio of expression was represented. (E) Expression and localization of TJP on HCT116 cells overexpressing Akt1 or Akt2 was checked by IF; Nucleus (Blue, DAPI), Actin peripheral cytoskeleton (Red, Phalloidin) and ZO1, top, or Claudin2, bottom, (Green).

**Supplementary figure 2.**
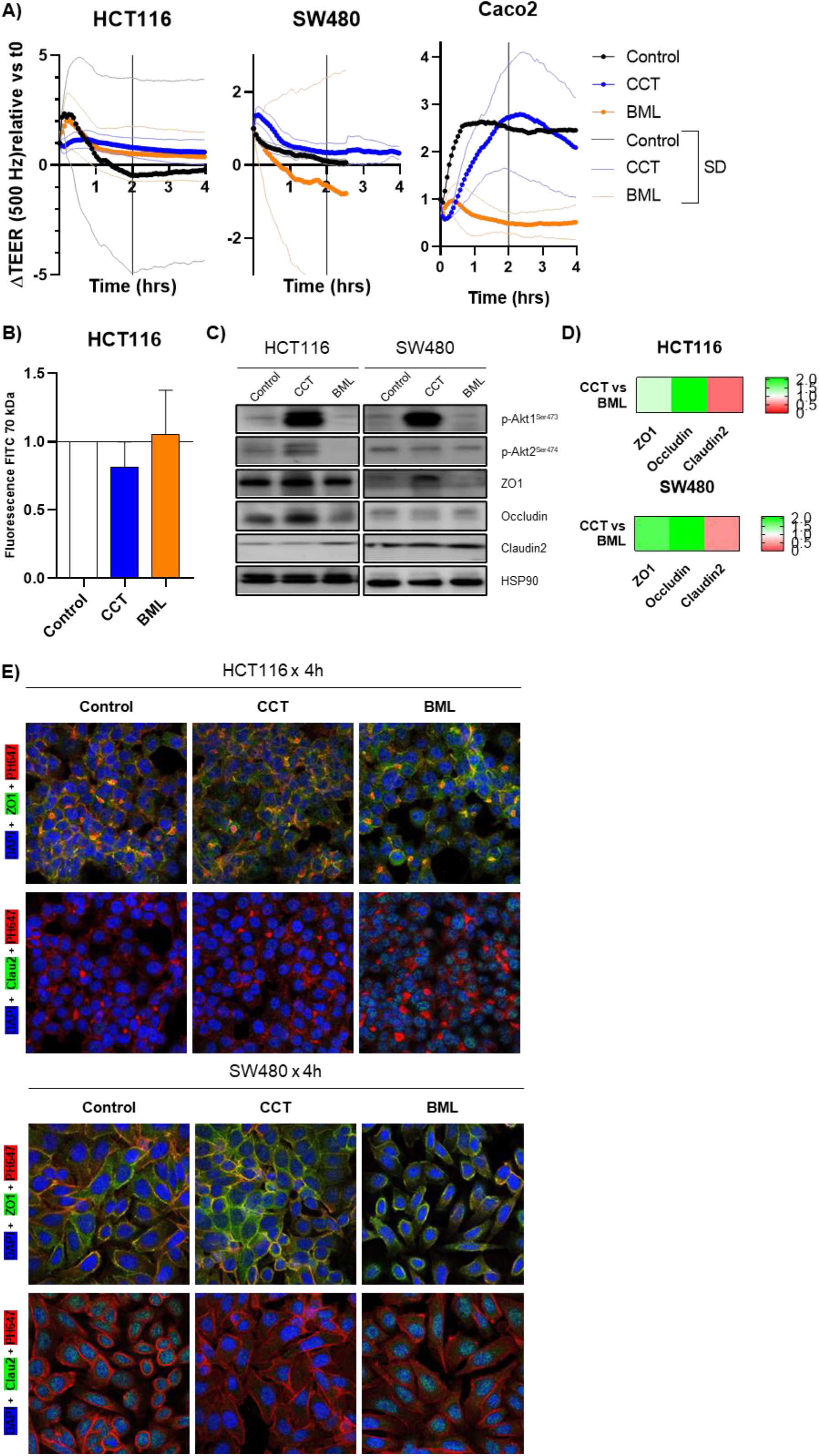
Pharmacological inhibition of Akt2 increases intestinal epithelial barrier function. (A) TEER assay was performed in HCT116, SW480 and Caco2 cells to determine the barrier membrane resistances of cells treated with the Akt inhibitors. (B) Permeability assay with FITC-Dextran 70 KDa 1h post FITC incubation on HCT116. Confluent HCT116 and SW480 cells were treated with Akt1 or Akt2 inhibitors for 30 minutes and TJPs and Akt phosphorylation were checked by (C) western blot and (D) quantification of 3 WBs; each band density was corrected by HSP90 expression. Then CCT/BML ratio of expression was represented. (E) IF staining of ZO1 and Claudin 2 in HCT116 and SW480 cells after treatment with Akt inhibitors, marking nucleus (Blue, DAPI), Actin peripheral cytoskeleton (Red, Phalloidin) and ZO1, top, or Claudin2, bottom, (Green).

**Supplementary Figure 3.**
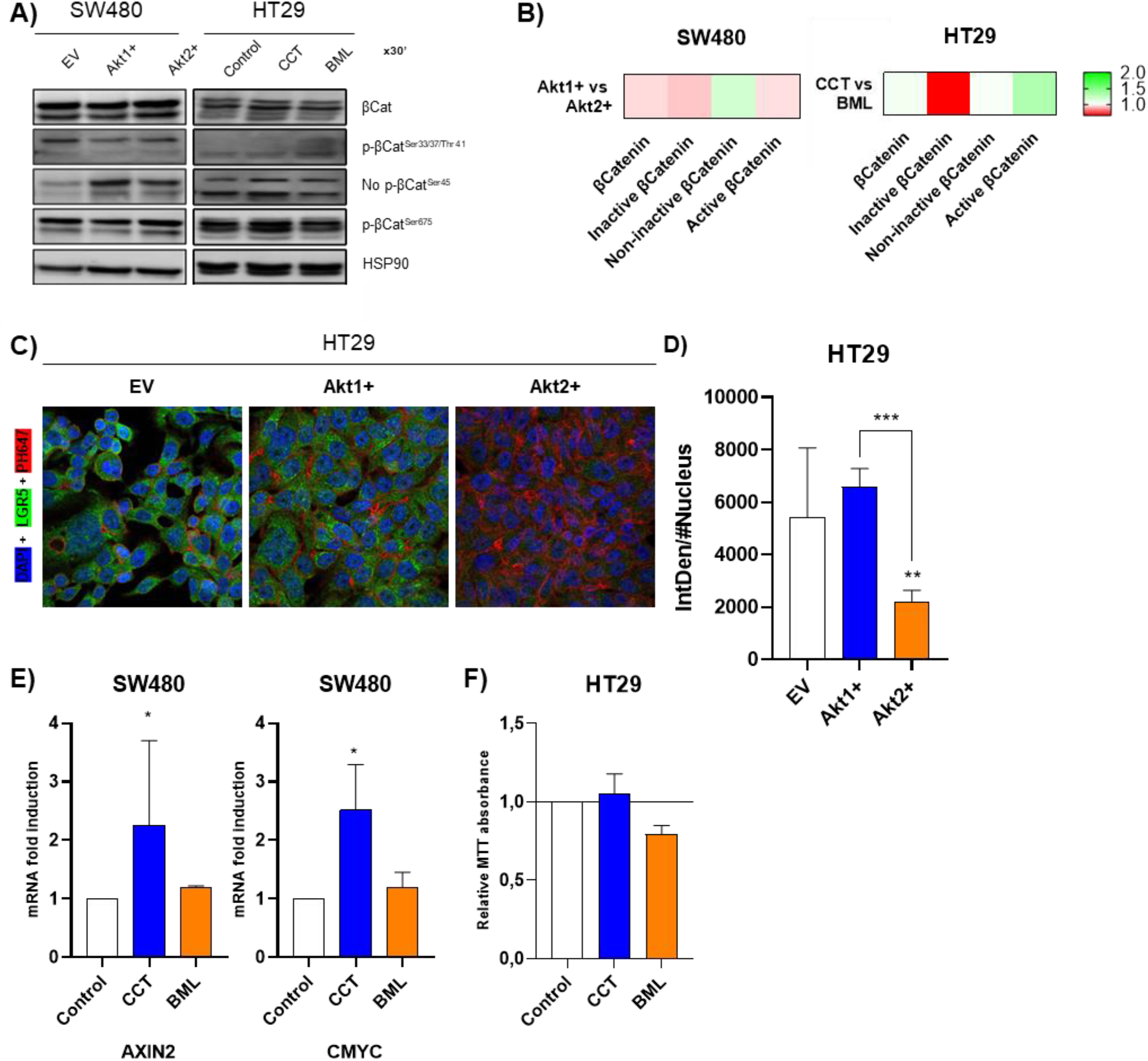
Akt inhibition controls β-Catenin phosphorylation and WNT pathway (A) Analysis of the state of activated (p-βCat Ser675 and No p-βCat Ser45) or marked to degradation (p-βCatSer33/37/Thr41) β-Catenin by western blot on confluent SW480 overexpressing cells and HT29 confluent cells treated with the inhibitors for 30 minutes and (B) quantification of 3 WBs; each band density was corrected by HSP90 expression. Then Akt1+/Akt2+ ratio of expression was represented. (C) HT29 overexpressing cells IF staining of LGR5, marking nucleus (Blue, DAPI), Actin peripheral cytoskeleton (Red, Phalloidin) and LGR5 (Green) and (D) IF quantification LGR5 intensity corrected by number of cells (Nucleus). (E) qPCR was performed on SW480 cells to check the changes on transcription of the targets of βCat/WNT pathway after β-Catenin translocation and TF activity; *Axin2* and *cMyc* mRNA levels after treatments with inhibitors for 24h and (E) MTT assay on HT29 IECs with the same conditions to check cell viability, survival and cell growth.

**Figure Supplementary 6.**
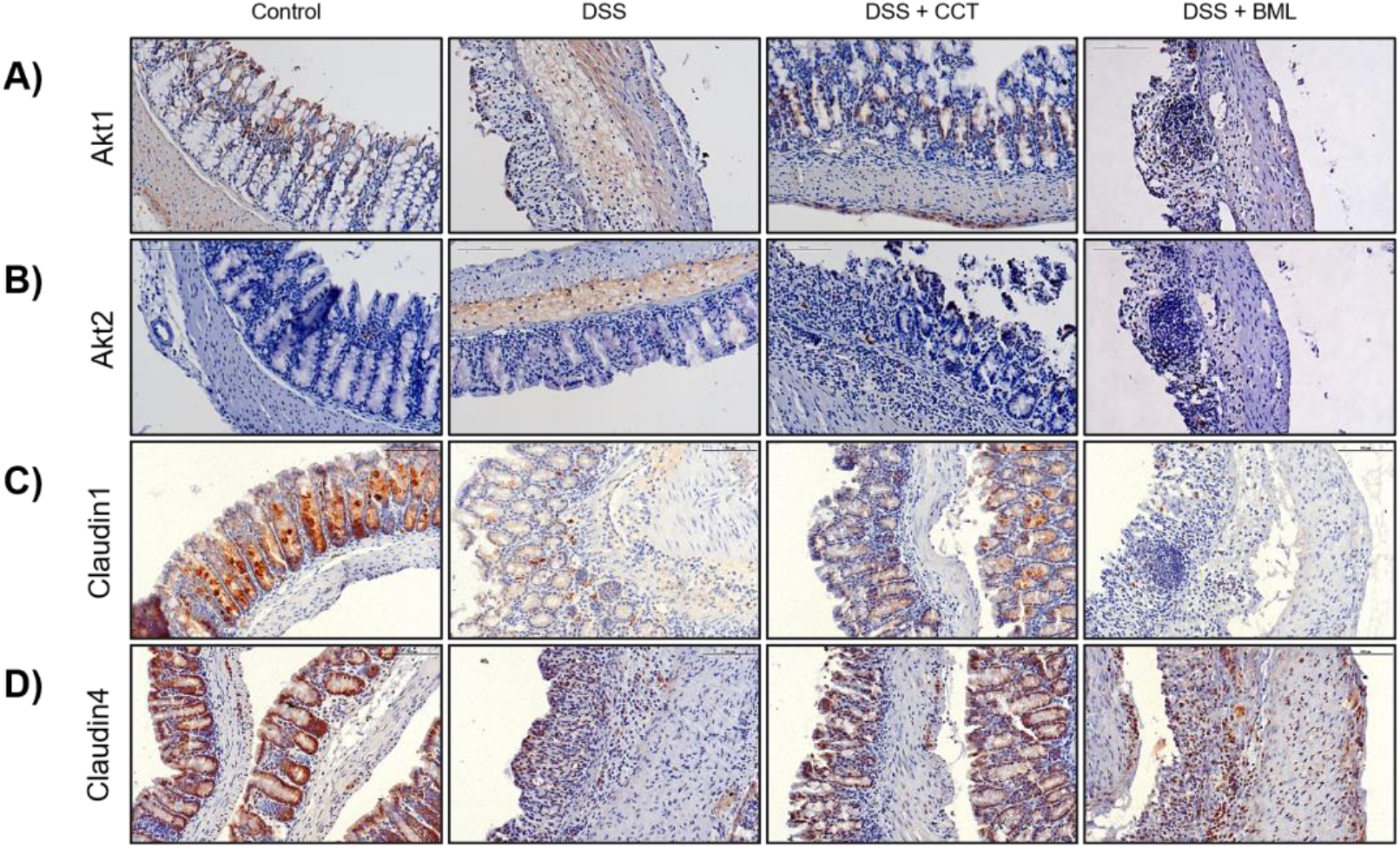
Immunohistologycal staining for (A) Akt1 and (B) Akt2 to check the effect of the inhibitors in situ on the colon tissue and for (C) Claudin1 and (D) Claudin4 as pro-barrier tight junctions’ proteins. Mice were treated with DSS 3% on drinking water ad libitum (DSS, disease control mice) and treated with either Akt2 inhibitor, CCT128 (DSS + CCT) or Akt1 inhibitor, BML-257 (DSS + BML), both at 1 mg/mouse, each day for a week. Control mice were given control water and vehicle treatment

